# Base Editor-Mediated Large-Scale Screening of Functional Mutations in Bacteria for Industrial Phenotypes

**DOI:** 10.1101/2022.09.27.509808

**Authors:** Yaomeng Yuan, Xihao Liao, Shuang Li, Xin-hui Xing, Chong Zhang

## Abstract

Base editing, the targeted introduction of point mutations into cellular DNA, holds promise for improving genome-scale functional genome screening to single-nucleotide resolution. Current efforts in prokaryotes, however, remain confined to loss-of-function screens using the premature stop codons-mediated gene inactivation library, which falls far short of fully releasing the potential of base editors. Here, we developed a base editor-mediated functional single nucleotide variant screening pipeline in *E. coli*. We constructed a library with 31,123 sgRNAs targeting 462 stress response-related genes in *E. coli*, and screened for adaptive mutations under isobutanol and furfural selective conditions. Guided by the screening results, we successfully identified several known and novel functional mutations. Our pipeline might be expanded to the optimization of other phenotypes or the strain engineering in other microorganisms.

## 1. Introduction

Microbial cell factories, as core components of green biomanufacturing and the bioeconomy, are widely applied in the bioproduction of fuels, medicines, chemicals and foods. Reverse engineering strategies, such as adaptive laboratory evolution (ALE) and random mutagenesis, are broadly employed to optimize the performance of microbial cell factories in the genome scale. However, these two methods suffer from a limitation in that only a small number of mutations could be detected within practical timescales, resulting in a loss of information^1^. Additionally, many engineered strains obtained by these strategies usually carry large amounts of mutations, making it laborious to identify those that play key roles in phenotype acquisition. Consequently, these methods are not efficient in providing useful knowledge to guide strain engineering in the future. To fully exploit bacterial genome resources and provide insights to microbiology and the engineering of microbial cell factories, it is crucial to develop high-throughput methods to reveal how genotypes give rise to phenotypes.

Forward genetic screening strategies, which introduce perturbations in the genome and characterize the corresponding phenotypes, have facilitated the acquisition of genotype-phenotype association (GPA) knowledge. Two high-throughput methods have been established to map GPA at single-gene resolution. The first relies on the transposon-based gene deletion library (Tn-seq), where GPA can be quantitatively identified by mixing transposon-insertion mutants and tracking their abundance under specific culture conditions via next-generation sequencing (NGS). Despite the great success, the random insertion of transposon hinders the application of Tn-seq to specific gene subsets. The establishment of CRISPR interference screening addresses the limitation, as genes of interest can be knocked down by designing sgRNA libraries^2–4^.However, these approaches are unable to reveal the role of specific positions within a gene. Moreover, mutations could be introduced to further modify the target genes identified by these tools for phenotype optimization. Recently developed CRISPR-Cas9-associated gene-editing tools, such as CREATE, have enabled the exploration of the effects of genomic loci at single-base resolution^5^. However, mutagenesis using CREATE relies on the formation of a double-strand break (DSB) followed by homology-directed repair in the presence of donor DNA, which often necessitates complex plasmid construction and strain background.

Base editors, where cytosine deaminase or adenine deaminase is fused to Cas9/Cas12 protein variants (e.g., dCas9, nCas9) to introduce specific mutations at target sites, have been shown to be effective in a variety of species^6–9^. Unlike CREATE, base editors create mutations by directly modifying the target bases without DSBs and recombination, making them easier to manipulate and applicable to a wide range of hosts. Base editor associated large-scale library screening has been applied in mammalian cells^10,11^ and *Saccharomyces cerevisiae*^12^. However, the procedures in these works were not suitable for prokaryotes due to the different genetic basis. Recently, the base editor mediated loss-of-function screening has been successfully applied in *Corynebacterium glutamicum*^13^, where stop codons were introduced to identify functional genes relevant to some specific traits. However, this study did not take full advantage of the ability of base editors to create additional types of amino acid substitutions and define functional mutations at base resolution.

Here, we exploited the potential of base editor-mediated pooled screening for functional single nucleotide variants (SNVs) in prokaryotes. We developed a dual-inducible Target-AID system for pooled screening and tested its ability to systematically modify genes by creating base substitutions at 31,123 individual sites across 462 stress response-associated genes in *E. coli*. As a proof of concept, two example screenings were performed to identify adaptive mutations under isobutanol and furfural selection pressure. As a result, several mutants exhibited 1.12∼1.45-fold enhanced growth in the selective conditions, demonstrating that our pipeline is a powerful tool for functional SNV defining and strain engineering.

## 2. Results

### 2.1 Establishing a dual-induced Target-AID version suitable for pooled screening

Base editor-mediated editing (Target-AID) has been shown to efficiently achieve point mutagenesis at target sites in *E. coli*^8^. In Target-AID, the *Petromyzon marinus* cytosine deaminase PmCDA119 (termed CDA), uracil DNA glycosylase inhibitor (termed UGI) from bacteriophage PBS2^17^ and protein degradation tag (LVA tag)^18^ were fused with the catalytically inactivated Cas9 variant (dCas9) from *Streptococcus pyogenes*. Guided by sgRNA, the dCas9-CDA-UGI-LVA fusion protein can introduce a cytosine to thymine conversion at the -14 to -20 positions from the PAM, with the highest efficiency observed at positions -17 ∼ -19^8^. However, the expression of Target-AID can also block gene transcription initiation and elongation when dCas9 binds to the target site in bacteria, resulting in decreased gene transcription levels (known as the CRISPRi effect). To ensure that fitness benefits of the variants are due to base substitution rather than gene knockdown, the expression of Target-AID should be turned off once mutagenesis is completed when applied in pooled screening. Therefore, we designed a dual-induced Target-AID system where the expression of dCas9-CDA-UGI-LVA protein was controlled by arabinose and the transcription of the corresponding sgRNA was regulated by rhamnose (Figure S1). After inducing mutagenesis, cells were washed and inoculated into the selection medium without inducers for screening (Figure 1A).

**Figure 1.**
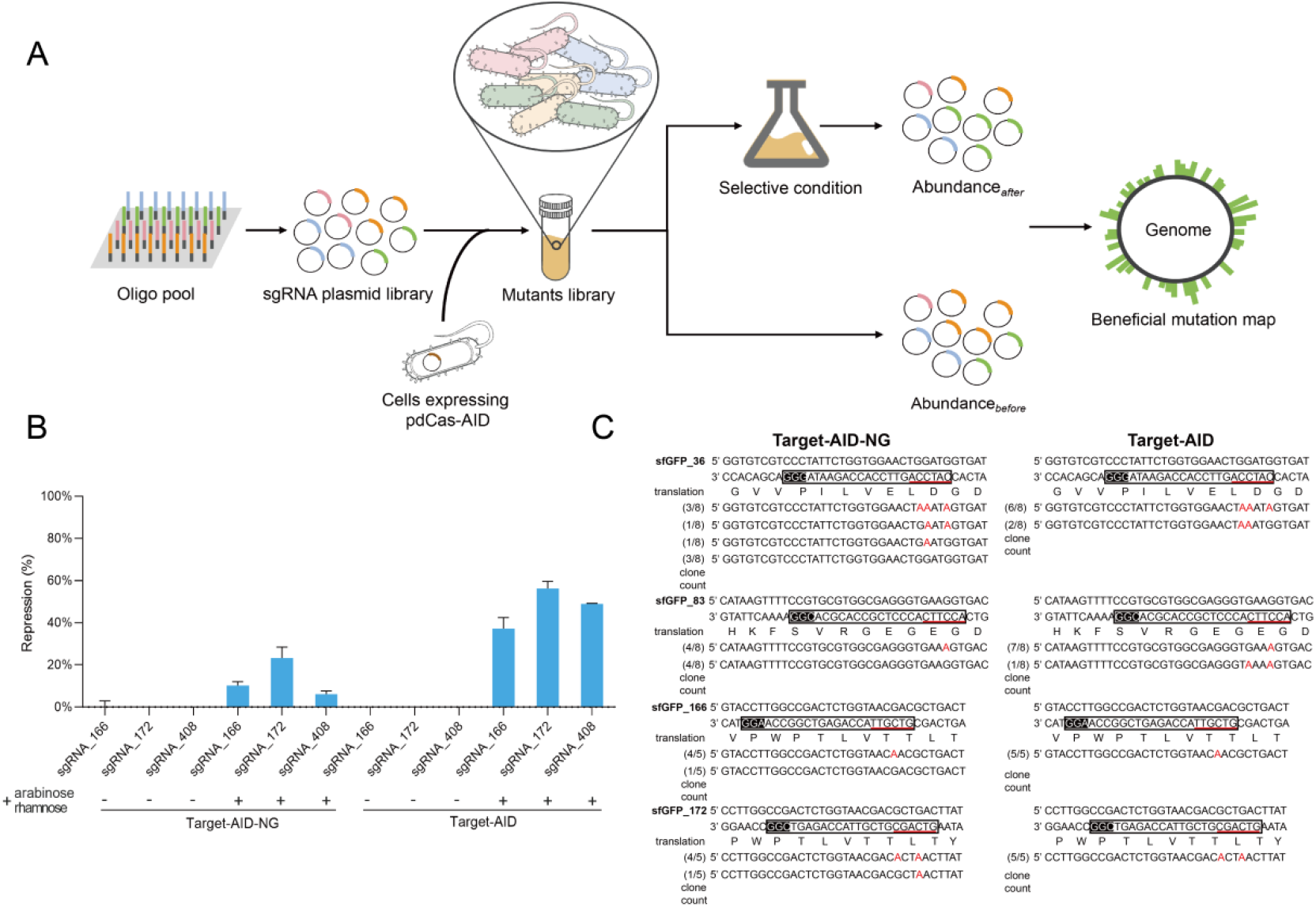
Target-AID mediated pooled screening workflow in *E. coli*. (A) Target-AID-mediated pooled screening workflow. (B) Characterization of fluorescence repression by Target-AID and Target-AID_NG. Target-AID indicates the base editor constructed using dCas9; Target-AID_NG indicates the base editor constructed using spCas9-NG. (C) Editing efficiency evaluation for Target-AID_NG and Target-AID by Sanger sequencing. sgRNA_36 and 83 were chosen to create missense mutations, and sgRNA_166 and172 were chosen to create synonymous mutations. The sgRNA targeting region and the PAM are framed in white and black boxes, respectively. The editing window is underlined in red, and the mutated bases are marked by red characters.

To evaluate the CRISPRi effect of the system, we used *sfGFP* as a reporter gene and introduced three synonymous mutations in its ORF via Target-AID. To expand the range of editable sites when designing the sgRNA library, a Cas9 protein variant, spCas9-NG^19^, with an expanded PAM of NG, was also used to construct the base editor (termed Target-AID-NG). In the absence of inducers, neither system exhibited significant fluorescence repression, indicating negligible CRISPRi effect or expression leakage. Upon induction, Target-AID system showed greater fluorescence repression, suggesting a stronger binding activity with target DNA (Figure 1B). We further evaluated the editing efficiency of both systems through Sanger sequencing of single colonies post-induction. Target-AID exhibited a higher mutation rate (Figure 1C) and was subsequently employed in our screenings.

### 2.2 Design and creation of mutagenesis library

We selected 462 genes associated with the stress response as targets^2,5,20^. Utilizing the principle previously established by our group, we searched for sgRNAs with high targeting specificity ^2^ and selected those containing cytosine(s) at positions -14 ∼ -20. Together with 400 non-targeting sgRNAs, our mutagenesis library comprised 31,523 sgRNAs, predicted to generate approximately 6,534 synonymous, 4,083 nonsense, and 33,476 missense mutations (Data S1, S2).

In this study, we chose isobutanol and furfural as selection pressures. Isobutanol, along with some other higher alcohols, is considered to be the most promising substitute for gasoline^21^. The toxicity of these biofuels to strains has been regarded as a key factor limiting their bioproduction^22^. Furfural is a major inhibitor of microbial fermentation as it is a toxic byproduct produced at high concentrations during lignocellulose pretreatment^23^. It impairs microbial growth by consuming NAD(P)H and inhibiting glycolysis and the TCA cycle^24^.

### 2.3 Target-AID mediated high-throughput functional screening

We applied the library to identify adaptive mutations in the selective environments. The library was transformed into *E. coli* BW25113 with 400-fold coverage and the mutant library was obtained by adding arabinose and rhamnose. We estimated editing efficiency by randomly picking ∼50 colonies and sequencing both sgRNAs and the corresponding genomic sites, obtaining an 82.5±0.71% mutation rate (Table S1). The variant library was then screened in M9 medium containing 4 g/L isobutanol or 0.5 g/L furfural, concentrations determined to significantly decrease cell growth in wild-type strain harboring non-targeting sgRNA (Figure S2). Two biological replicates were set up for each screening. For NGS data analysis, we filtered out the low-abundance sgRNAs in the synthesized plasmid library and calculated the fitness of remaining sgRNAs as described in the Methods. Pearson coefficients of sgRNA fitness between two replicates were higher than 0.9 in both screenings, indicating high consistency (Table S2). 255 and 386 sgRNAs were significantly enriched in furfural and isobutanol screenings, respectively (FDR<0.05).

When comparing the top 100 enriched sgRNA in both screenings, we observed several shared sgRNAs (Figure 2). Notably, A large proportion of sgRNAs targeting *pcnB* showed high fitness in both screenings. We hypothesize that these shared mutations may enhance the growth in multiple selection environments. We also observed some unique patterns in different screenings. In furfural selection, *GcvA*, encoding a dual-regulator of glycine cleavage system, had 7 enriched sgRNAs in the top 100 hits, with 5 creating premature stop codons. In isobutanol selection, multiple mutations in the RNA polymerase subunit β (encoded by *rpoB*) were highly enriched.

**Figure 2.**
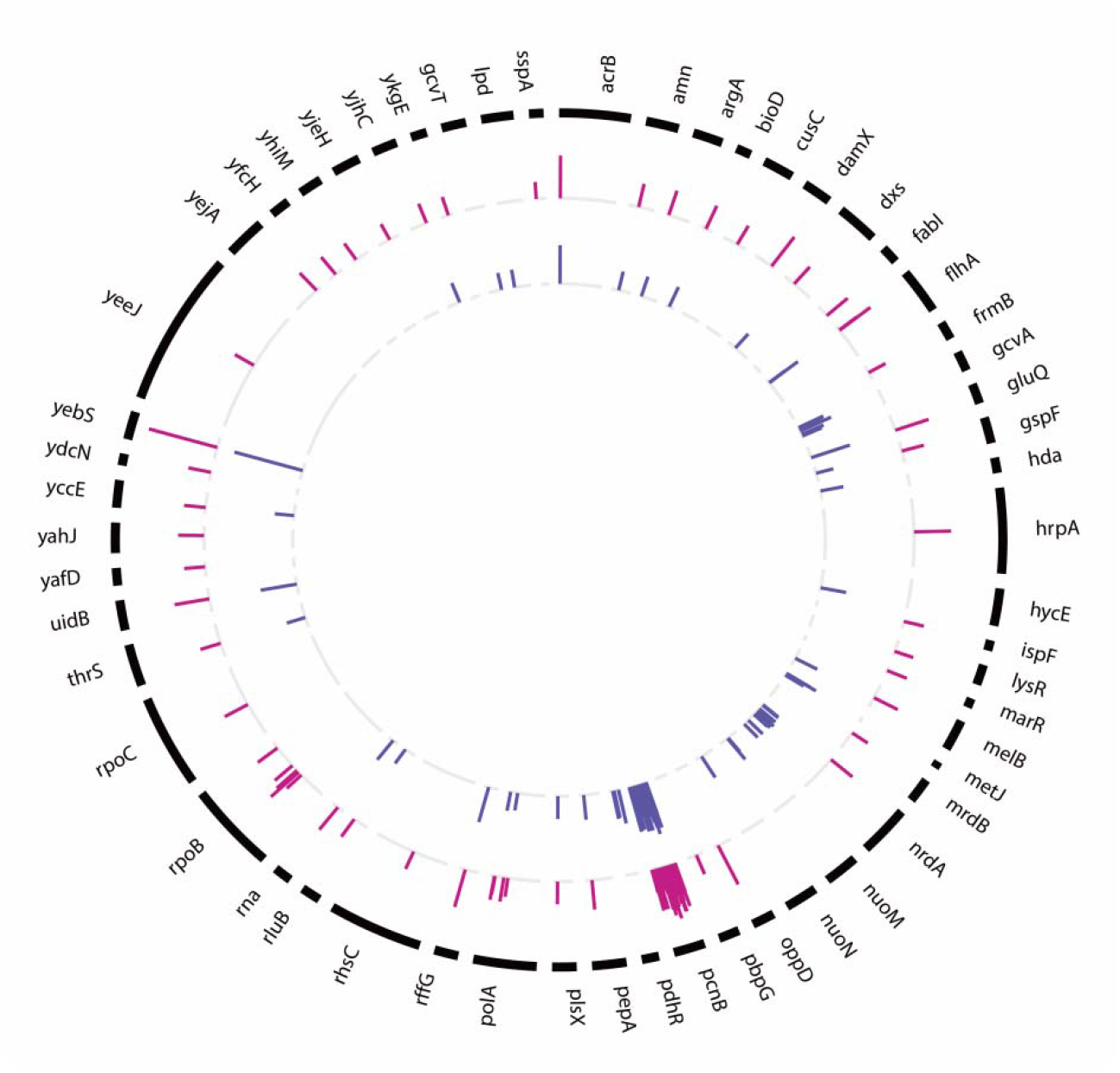
Fitness of the top 100 enriched sgRNAs (FDR<0.05) and the corresponding target genes under different conditions. From the outer circle to inner circle, the length of the bars represents the fitness of sgRNAs in the presence of isobutanol and furfural, respectively.

### 3.4 Mutants in *pcnB* improved the growth by lower the plasmid copy number

To confirm our population-level analysis, we reconstructed the top 10 most enriched variants and verified their growth curves individually (Figure 3A, 3B). Nine out of ten mutations were located in *pcnB* gene in both screenings (Table 1). In the presence of isobutanol, the best reconstructed mutants, *pcnB*_R203H_E204K_D205N, *pcnB*_G103D_R104H and *pcnB*_G183N, showed 1.45-, 1.37- and 1.31-fold higher growth than the control strain, respectively (Figure 3C). When testing the growth of reconstructed strains in the presence of furfural, the *pcnB*_R203H_E204K_D205N and *pcnB*_G103D_R104H variants again exhibited the greatest growth improvement (1.12- and 1.15-fold, Figure 3D).

**Table 1.**
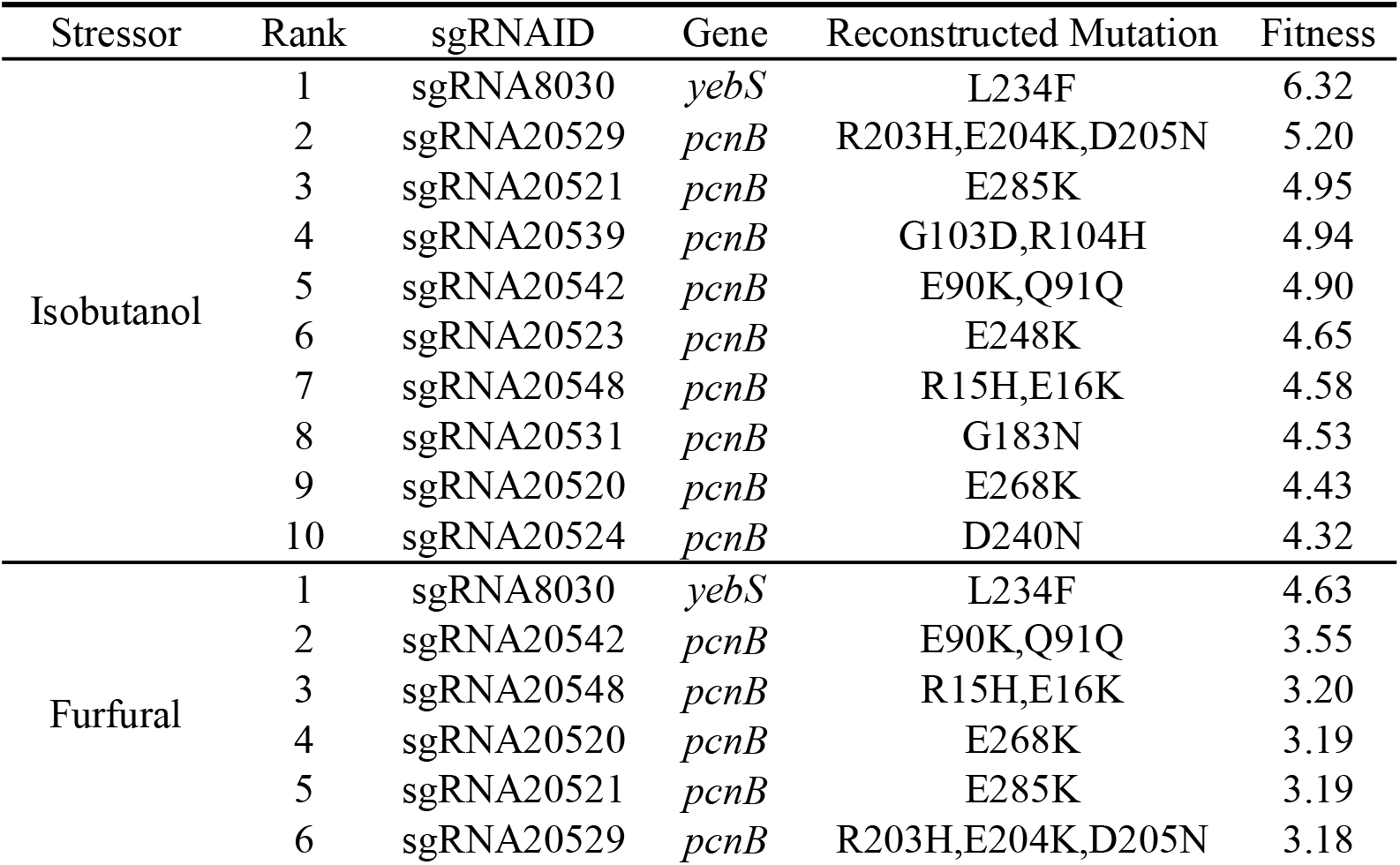
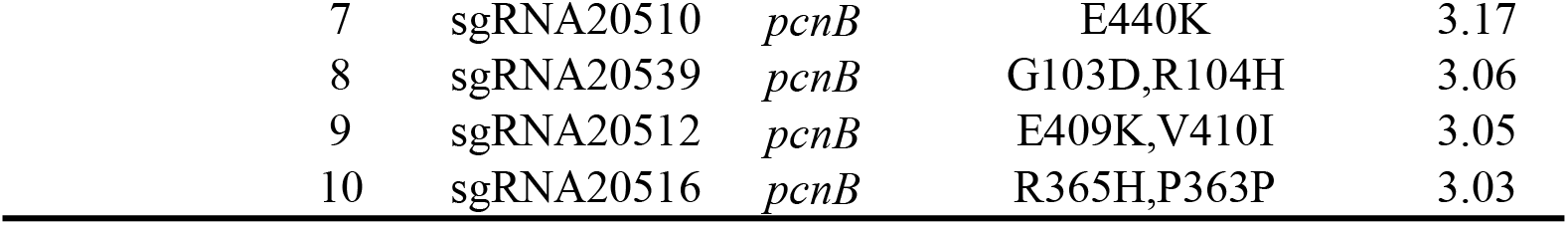
Top 10 enriched mutations in each screening (FDR<0.05)

| Stressor | Rank | sgRNAID | Gene | Reconstructed Mutation | Fitness |
| --- | --- | --- | --- | --- | --- |
| Isobutanol | 1 | sgRNA8030 | *yebS* | L234F | 6.32 |
|  | 2 | sgRNA20529 | *pcnB* | R203H,E204K,D205N | 5.20 |
|  | 3 | sgRNA20521 | *pcnB* | E285K | 4.95 |
|  | 4 | sgRNA20539 | *pcnB* | G103D,R104H | 4.94 |
|  | 5 | sgRNA20542 | *pcnB* | E90K,Q91Q | 4.90 |
|  | 6 | sgRNA20523 | *pcnB* | E248K | 4.65 |
|  | 7 | sgRNA20548 | *pcnB* | R15H,E16K | 4.58 |
|  | 8 | sgRNA20531 | *pcnB* | G183N | 4.53 |
|  | 9 | sgRNA20520 | *pcnB* | E268K | 4.43 |
|  | 10 | sgRNA20524 | *pcnB* | D240N | 4.32 |
| Furfural | 1 | sgRNA8030 | *yebS* | L234F | 4.63 |
|  | 2 | sgRNA20542 | *pcnB* | E90K,Q91Q | 3.55 |
|  | 3 | sgRNA20548 | *pcnB* | R15H,E16K | 3.20 |
|  | 4 | sgRNA20520 | *pcnB* | E268K | 3.19 |
|  | 5 | sgRNA20521 | *pcnB* | E285K | 3.19 |
|  | 6 | sgRNA20529 | *pcnB* | R203H,E204K,D205N | 3.18 |

|  |  |  |  |  |
| --- | --- | --- | --- | --- |
| 7 | sgRNA20510 | *pcnB* | E440K | 3.17 |
| 8 | sgRNA20539 | *pcnB* | G103D,R104H | 3.06 |
| 9 | sgRNA20512 | *pcnB* | E409K,V410I | 3.05 |
| 10 | sgRNA20516 | *pcnB* | R365H,P363P | 3.03 |

**Figure 3.**
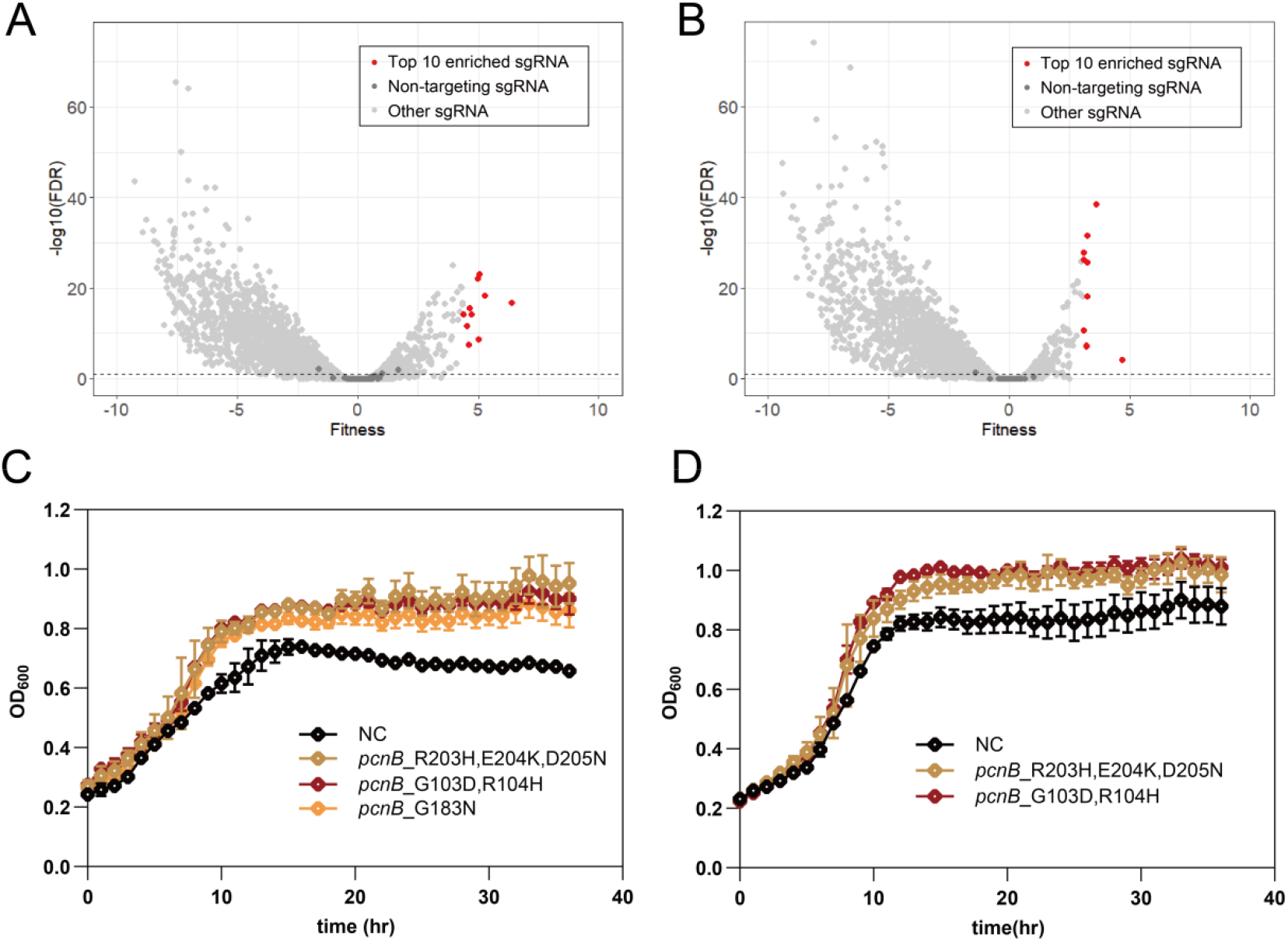
Population-level analysis results and growth validation in two selective environments. (A) Volcano plot of sgRNA fitness relative to -Log10(FDR) in the presence of isobutanol. (B) Volcano plot of sgRNA fitness relative to -Log10(FDR) in the presence of furfural. (C) Growth verification of top 10 enriched mutants in the presence of 4 g/L isobutanol. Variants with the most significant growth improvement were shown. The growth curves of the remaining reconstructed strains were shown in Figure S3. NC indicates the strain harboring non-targeting sgRNA. Results represent mean values and standard deviation of biological triplicates (N=3). (D) Growth verification of top 10 enriched mutants in the presence of furfural.

*pcnB* encodes poly(A) polymerase I, which adds polyA tails onto the 3’ end of RNA using ATP as a substrate. Poly(A) polymerase I plays a significant role in the global gene expression control by influencing the stability of *E. coli* transcriptome^25^. Knockdown or mutations of this gene have been observed to contribute to tolerance under stressed conditions, leading to suggestions that pcnB is a potential global regulator of stress tolerance ^2,26^. However, we noted that this gene is also involved in plasmid copy number control. In *pcnB-*deleted strains, the copy number of ColE-derived plasmids decreases due to increased stability of the RNase E processed form of RNA1, the antisense RNA regulator of ColE1 replication^27^. The sgRNA plasmid used in this study has a pMB1 origin of replication (ORI) belonging to the ColE family. Similarly, strains used in the studies where *pcnB* was found to increase tolerance carried ColE-derived plasmids. Therefore, the improved growth of *pcnB* mutants is most likely due to the decreased copy number of plasmids which lowered the metabolic burden.

To demonstrate our inference, we introduced the *pcnB*_R203H_E204K_D205N, *pcnB*_G103D_R104H and *pcnB*_G183N mutations in the genome of *E. coli* BW25113 using the curable plasmid-mediated CRISPR-Cas9 recombineering to construct 3 plasmid-free mutants (termed as P1, P2 and P3). However, we failed to construct the P2 strain due to the unexpected frameshift mutations. For the remaining 2 strains, neither of them showed improved growth in the presence of isobutanol or furfural (Figure S4). To further validate the plasmid copy number was reduced in *pcnB* mutants, we constructed a test plasmid pRFP by inserting an RFP expression cassette into the sgRNA plasmid backbone. The resulting pRFP was then chemically transformed into P1, P3, and wild-type strains. P1 and P3 exhibited significantly reduced fluorescence intensity (Figure 4A) and the growth improvement was recovered (Figure 4B, 4C). These results suggest that the enhanced growth of *pcnB* mutants was due to reduced plasmid copy number. R203 has been proven to cause a more than 90% decreased activity of poly(A) polymerase I when mutated into alanine in a previous study^28^. We propose that the mutations identified here are function disruptive ones.

**Figure 4.**
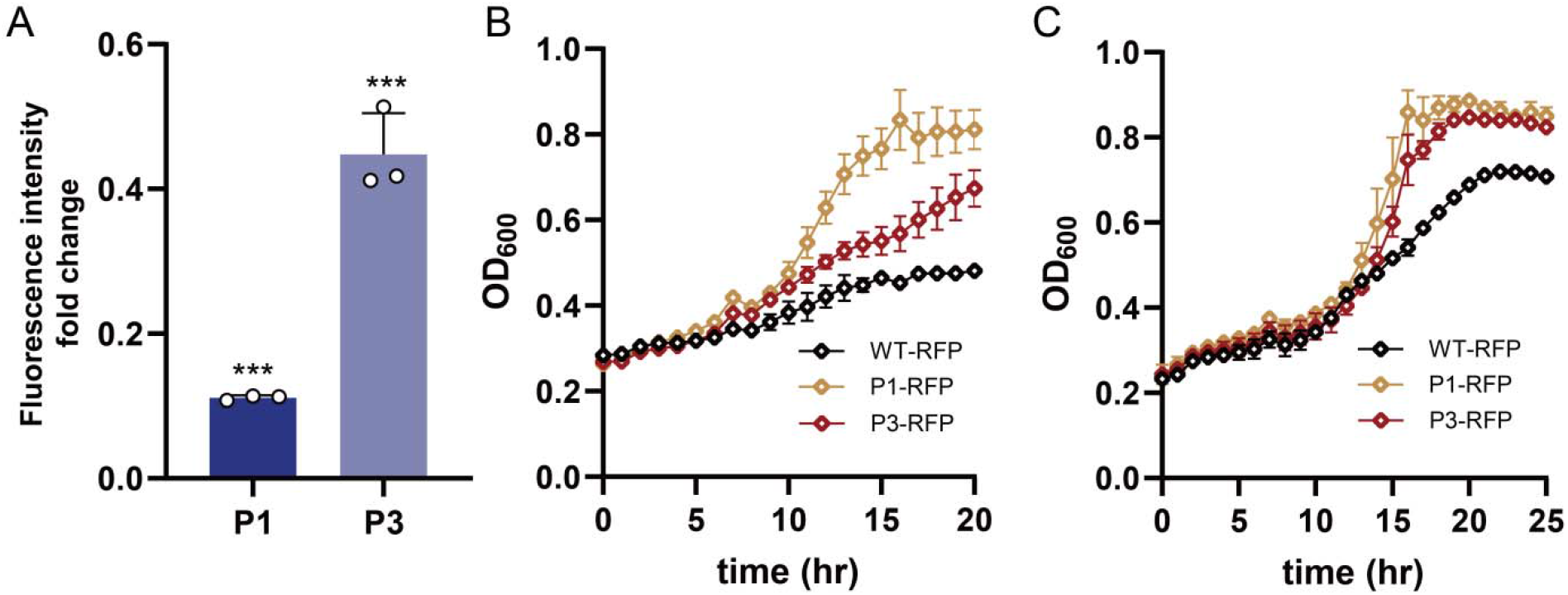
Plasmid copy number evaluation in *pcnB* mutants. (A) Fluorescence intensity fold change of P1 and P3 compared with wild-type strain. Results represent mean values and standard deviation of biological triplicates (N=3). Two tailed t-test were performed to determine the significance levels against the wild-type strain. ***P<0.001. (B) Growth curves of P1, P3 and wild-type strain harboring pRFP in M9 medium containing 5 g/L isobutanol. Results represent mean values and standard deviation of biological triplicates (N=3). (C) Growth curves of P1, P3 and wild-type strain harboring pRFP in M9 medium containing 1 g/L furfural.

### 3.5 *GcvA* inactivation showed increased fitness in the presence of furfural

We then reconstructed the top 3 most enriched *gcvA* inactivation variants (*gcvA*_W108*, *gcvA*_Q201*, *gcvA*_Q245*). All of the mutants showed an increased growth rate compared with the control strain in the presence of furfural (Figure 5A). Since all 3 *gcvA* mutants harbored nonsense mutations, we further knocked out *gcvA* in *E. coli* BW25113 and found that the mutant exhibited a faster growth compared to the wild-type strain (Figure 5B). Glycine cleavage system, encoded by *gcvTHP* operon, cleaves glycine to produce 5,10-methylenetetrahydrofolate (5,10-mTHF) required for serine, methionine, thymidine, and purine biosynthesis. Expression of the *gcv* operon is activated by *gcvA* in the presence of glycine and repressed when glycine is limiting. A previous study has shown that deletion of *gcvA* results in reduced expression of the *gcv* operon^29^. Another research observed that the expression level of *gcvH* and *gcvP* was down-regulated under multiple stressors, indicating many-sided roles of *gcvTHP* beyond amino acid metabolism^30^. *GcvA* has also been reported as a repressor of the *hdeAB* operon which confers higher acid tolerance by protecting periplasmic proteins from aggregation^31^. We speculate the deletion of *gcvA* improved growth phenotype by modifying multiple cellular processes.

**Figure 5.**
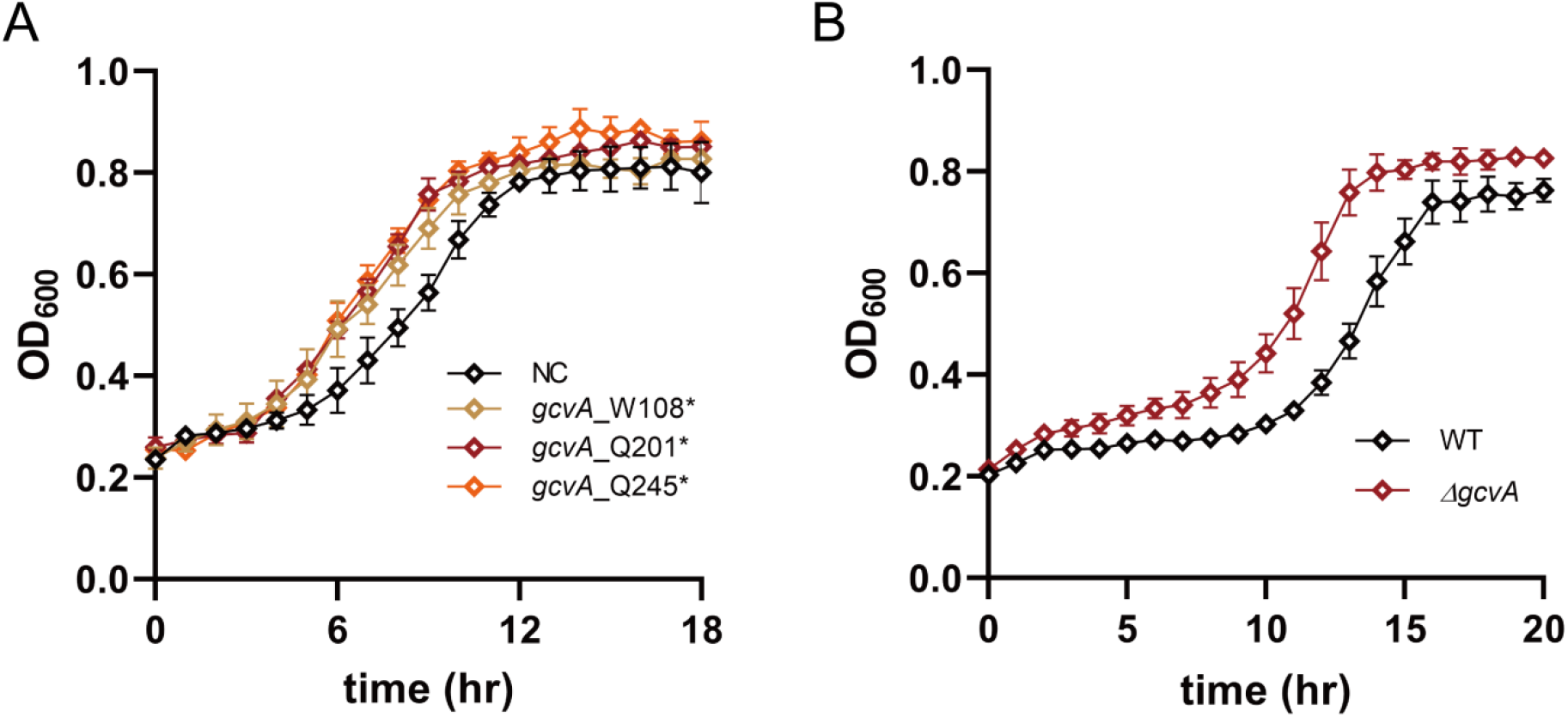
Growth curves of the *gcvA* inactivation variants. (A) Growth verification of gcvA mutants in the presence of 0.5 g/L furfural. Mutants were constructed using Target-AID. NC indicates the strain harboring non-targeting sgRNA. Results represent mean values and standard deviation of biological triplicates (N=3). (B) Growth verification of gcvA-deleted strain in the presence of 1 g/L furfural. Gene deletion was achieved by the plasmid-curable CRISPR-Cas9 recombineering. WT indicates wild-type strain. Results represent mean values and standard deviation of biological triplicates (N=3).

### 3.6 Mutations in *rpoB* enabled a faster growth in the presence of isobutanol

Mutations in *rpoB* have been repeatedly observed to confer a fitness benefit under various stressed conditions^32–34^. In our study, multiple *rpoB* mutants were highly enriched in isobutanol selection. Therefore, we reconstructed the top 5 enriched mutations in *rpoB* (*rpoB*_S621F,N620N, *rpoB*_A665V, *rpoB*_S788F, *rpoB*_G1218N,E1219K and *rpoB*_E546K) and tested the growth curves individually. Mutations rpoB_S788F and *rpoB*_S621F,N620N enabled a 43% and 25% faster growth rate in the glucose consumption phase (Figure 6), while *rpoB*_E546K showed a slightly higher growth (Figure S6). The remaining two mutants exhibited little growth improvement (Figure S5). Previous studies have shown that variants at positions E546X and E672X resulted in similar growth improvement phenotypes in several selection environments because part of the maintenance energy shifted to growth-related processes^35^. Since the beneficial *rpoB* mutants identified here are located in the same structure community as E546 and E672^35^, they may regulate the transcriptome in a similar way. Further research is needed to explain the underlying molecular mechanism.

**Figure 6.**
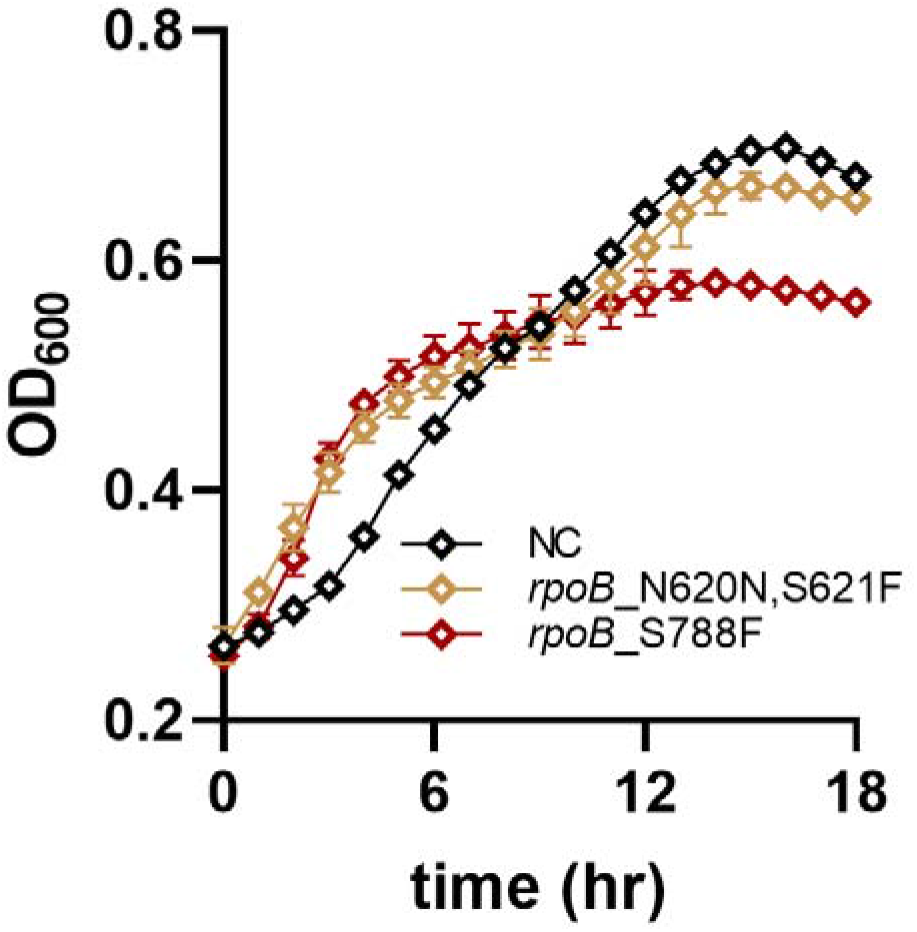
Growth validation of strain with *rpoB*_S788F and *rpoB*_S621F,N620N mutations. NC indicates the strain harboring non-targeting sgRNA. Results represent mean values and standard deviation of biological triplicates (N=3).

## 3. Discussion

Large-scale functional screening mediated by the tools derived from CRISPR-Cas9 systems accelerated the mapping of genotype-phenotype associations in a trackable and customized manner. Although CRISPRi pooled screening has been widely applied in functional genomic studies, it can only identify how single genes contribute to the phenotype. At the single base level, CREATE is a representative approach for precise genome editing in prokaryotes, where base substitution is generated by homology-directed repair of DSBs in the presence of donor DNA. Despite great success, the toxicity of DSBs, the requirement of DNA templates and the complex strain background hinder its broader application. In contrast, base editors enable precise gene editing that is not limited by the aforementioned conditions. To fully harness the power of base editors in screening functional loci at single-base resolution in prokaryotes, we proposed a Target-AID mediated large-scale library screening workflow in this work. To demonstrate the feasibility of our pipeline, we constructed an sgRNA library that could introduce mutations at 31,123 individual sites across 462 genes and applied it to search beneficial mutations under stressors. Known and novel adaptive mutations were successfully identified, demonstrating the capability of our strategy in engineering and discriminating functional variants.

We speculate that some functional mutations cannot be enriched due to the low editing efficiency of some sgRNAs. Moreover, one sgRNA may create several kinds of mutations within the editing window. Different effects of these mutations on growth may interfere with sgRNA fitness. For example, a mixture of neutral and beneficial mutations may lead to underestimation of sgRNA fitness. On the other hand, although some sgRNAs showed increased fitness in the screening, the growth improvement of the corresponding variants may be too insignificant to be detect in validation. In addition, based on the screening data analysis and growth verification, we found that some enriched mutants with low initial read counts showed no significant growth improvement (i.e., *yebS* mutant, Figure S4), which was also observed by Liu et al.^13^ For more accurate and efficient genotype-phenotype association, more research should be performed to improve base editor performance and data analysis algorithms.

Recently, base editors with improved performance have been reported, paving the way for the development of functional screening mediated by base editors. Developing base editors with a broad PAM by using new Cas9 variants can widen the editable scope in the genome. In this work, we also tested the performance of the Target-AID-NG constructed by using SpCas9-NG^36^. However, this base editor was not chosen to conduct pooled screening due to the lower editing efficiency. How to maintain high editing activity while widening the PAM site is an important problem to be solved in the future. Establishing base editors with fewer bystanders, higher editing efficiency, and reduced off-target activity and developing algorithms to predict editing outcomes will also improve predictability and reduce the difficulty in library design and data analysis ^37–39^. Compared with CREATE, one of the major drawbacks of Target-AID is the limited base mutation type in *E. coli* (C to T). Recently, C to A conversion was observed in *E. coli* MG1655 when using Ung-nCas9-AID fusion^40^. Together with the development of adenine base editors (ABEs)^7^, exploiting the potential of these new base editors may further expand their capability in functional screenings. Furthermore, by combining biosensors or chromogenic reactions with screening platforms such as fluorescence-activated cell sorting (FACS)^41^, fluorescence-activated droplet sorting (FADS)^42^, droplet-based FACS^43^, and Gel-FACS^44^, more complex phenotypes such as metabolite production and protein secretion can also be screened. Finally, a curable plasmid can be designed for the base editing system to allow mutation combination via iterative screenings, which could potentially lead to better optimization of the desired phenotypes. Taken together, we are positive that base editor-mediated functional screening will accelerate large-scale interrogation of phenotype-genotype relationships in a broad range of hosts for metabolic engineering and synthetic biology.

## 4. Materials and Methods

### 4.1 Strain and plasmid construction

All strains, plasmids and primers used in this work are listed in Tables S3 and S4. *E. coli* BW25113 was obtained from ATCC. *E. coli* DH5α was purchased from Biomed and used for plasmid construction. *E. coli* s17-1 sfGFP was a gift of the Ceorge Guoqiang Chen laboratory at Tsinghua University.

Plasmids were constructed by Gibson assembly or GolgenGate assembly. The dCas9-CDA-UGI-LVA expression cassette was amplified from the pScI-dCas9-CDA-UL plasmid (Addgene #108551) and inserted into the pBAD43 vector via Gibson assembly to obtain pdCas9-AID. For pdCas9-AID-NG, the dCas9 sequence in pdCas9-AID was replaced by dspCas9-NG. The plasmid for sgRNA expression, termed pTarget, was derived from pTargetF_lac^2^ by replacing the lac promoter with the rhaBAD promoter. To facilitate library insertion into pTarget, pTarget_rhaBAD_RFP was further constructed by replacing the 20-bp target sequence with an RFP expression cassette containing two BsmBI sites in opposite orientations at both ends. A pair of oligonucleotide DNAs for construction of the targeting sgRNA vector was designed as follows: 5’-tagc-(target sequence)-3’ and 5’-aaac- (reverse complement of the target sequence)-3’. Goldengate assembly was performed to insert the target sequences into pTarget_rhaBAD_RFP to obtain pTarget series plasmids. All the constructed plasmids were confirmed by Sanger sequencing.

### 4.2 Assay of Target-AID mediated mutaganasis

*E. coli* BW25113 cells were chemically transformed with the appropriate plasmids and precultured at 37°C with 800 μL LB broth [10 g/L tryptone, 5 g/L yeast extract, 10 g/L NaCl] for 1 h. The cells were then transferred into 10 mL LB broth containing streptomycin (20 mg/L), ampicillin (100 mg/L), L-arabinose (100 mg/L) and L-rhamnose (300 mg/L) and grown overnight at 37°C and 220 r.p.m. The cultures were then serially diluted onto LB agar plates supplemented with appropriate antibiotics and incubated overnight at 37□. Mutations were confirmed by picking single colonies for Sanger sequencing.

### 4.3 Fluorescence characterization of the Target-AID system

For fluorescence characterization, *E. coli* s17-1 harboring the *sfGFP* gene was used as the host. Three sgRNAs targeting the *sfGFP* gene, as well as the non-targeting sgRNA, were chemically transformed into the s17-1 strain containing the pdCas9-AID or pdCas9-AID-NG plasmid. Subsequently, overnight cultures in LB medium from single colonies of these strains were individually incubated in 10□mL fresh LB broth in 50-mL flasks (initial OD_600_□=□0.02) with or without inducers for 12 h, and fluorescence was measured with an F-2500 Hitachi Fluorescence Reader (excitation, 488□nm; emission, 510□nm). Fluorescence was then normalized to the culture OD_600_ value measured on the Amersham Bioscience spectrophotometer. The repression ratio was calculated by comparing the relative fluorescence with respect to the control strain expressing the non-targeting sgRNA.

### 4.4 Design and preparation of the sgRNA library

462 target genes were chosen as editing targets, in which 35 genes were related to the resistance of acetate^5^, 97 genes were related to the tolerance of furfural and isobutanol^2^, and 330 genes were related to thermotolerance^14^. To ensure the activity and specificity of sgRNAs, we first selected all sgRNAs targeting the 462 genes with an off-target threshold of 15 according to our previous work^2^. Mutations were predicted according to the previously reported editing window and efficiency. The plasmid library was synthesized by Genewiz (termed ‘Plasmid Library’).

### 4.5 Screening experiments

The sgRNA library was electroporated into *E. coli* BW25113 carrying the pdCas9-AID plasmid. For competent cell preparation, *E. coli* BW25113 containing pdCas9-AID was grown in 100 mL LB medium without NaCl at 37 □ until the OD_600_ reached 0.6. The cells were then collected by centrifugation at 4 □, washed five times in ice-cold deionized water and resuspended in 5 mL 15% glycerol. The prepared competent cells were mixed with 500 ng library plasmid/mL competent cells, divided into 100 μL aliquots and added into 25-well electroporation plates. For electroporation, the BTX Harvard apparatus ECM 630 High Throughput Electroporation System was used, and the parameters were set as 2.1□kV, 1□kΩ, and 25□μF. The transformed cells were incubated in LB broth (1:4 v/v) for 1 h at 37°C and 220 r.p.m. Fifty microliters of the culture was diluted and streaked onto LB agar plates with streptomycin (20 mg/L) and ampicillin (100 mg/L) to calculate the electroporation efficiency and library coverage. The remaining culture was transferred into 100 mL LB medium containing streptomycin, ampicillin, L-arabinose (100 mg/L) and L-rhamnose (300 mg/L) and grown at 37 □ and 220 r.p.m. until the OD_600_ reached ∼1. We took 5 mL of the culture to extract plasmids and termed it ‘Before Library’ for NGS. The remaining cells were further collected by centrifugation, washed twice and resuspended in M9 medium (12.8 g Na_2_HPO_4_·7H_2_O, 3 g KH_2_PO_4_, 0.5 g NaCl, 1 g NH_4_Cl, 4 g glucose, 2 mM MgSO_4_, 0.1 mM CaCl_2_, for 1 L). Fifty microliters of the culture were diluted and spread onto LB agar plates, and 50 colonies were randomly picked for Sanger sequencing to calculate the mutagenesis efficiency. The remaining cells were inoculated into 100 mL M9 medium containing 4 g/L isobutanol or 0.5 g/L furfural with an initial OD_600_ of 0.03 and cultured until an OD_600_ of 1 was reached. 5 mL of the culture was taken for plasmid extraction and termed ‘After Library’. Two biological replicates were performed for each screening.

### 4.6 Next-generation sequencing (NGS) library sequencing and data analysis

The extracted plasmids were used as templates for PCR to amplify the N20 region of the sgRNAs. The PCR conditions were set as follows: one reaction (50 μL) per library, 50 ng templates per reaction, primer NGS-seq-F/R, KAPA HiFi HotStart polymerase (KAPA Biosystems), 95□□ for 3□min, 20 cycles (98□□, 20□s; 60□, 15□s; 72□□, 30□s), followed by 72□□ for 1□min. The amplified fragments were confirmed by 4% agarose gel electrophoresis, cleaned using a Gel Extraction Kit (Omega) and processed for NGS using standard Illumina preparation kits. Sequencing was carried out using a 2 ×□150 paired-end configuration.

The raw NGS data were first demultiplexed and filtered, and the adaptor was removed to obtain clean data for each library. Then, we merged each of the two pairs by FLASH script and removed the reads without detected pairs. Customized python scripts were applied to calculate the read counts of each sgRNA. To increase statistical robustness, we removed sgRNAs with <20 normalized read counts from the ‘Plasmid Library’. For the remaining sgRNAs, the read counts were processed by edgeR package^15^ and the Log_2_FC (Fold Change) was calculated using equations 1.

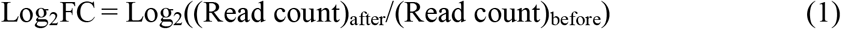

The fitness of each sgRNA was then calculated using equation 2, where NCsgRNA indicates non-targeting sgRNA.

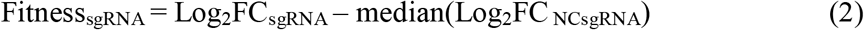

### 4.7 Mutant reconstruction and growth curve characterization

Mutations were introduced into *E. coli* BW25113 by base editor or the curable-plasmid mediated CRISPR-Cas9 recombineering. For base editing, mutations were created as described above. For gene deletion or mutation introducing by CRISPR-Cas9 recombineering, the experiments were performed as described by Jiang et al^16^. The sgRNAs were designed to target the corresponding sites (Table S1). Two homologous arms flanking at both sides of the target site were amplified separately using the primers containing mutated bases (Table S2). Overlap extension PCR was then applied to construct the donor DNA fragments. Mutations in the transformants were confirmed by colony PCR and Sanger sequencing. For the successfully edited colonies, plasmids were cured to obtain the plasmid-free strains.

The reconstructed strains were stored at -80°C. For growth curve verification, the stored strains were streaked onto LB plates to isolate single colonies. The colonies were then inoculated in LB broth for 6 h with shaking at 37°C and 220 r.p.m. Subsequently, the cells were collected, washed twice and inoculated into M9 medium containing the corresponding toxic chemicals to characterize growth using Bioscreen C MBR (Oy Growth Curves Ab Ltd). For the mutants reconstructed by base editor, 4 g/L isobutanol or 0.5 g/L furfural was added in the medium. For the mutants constructed by CRISPR-Cas9 recombineering, 5 g/L isobutanol or 1 g/L furfural, which inhibited the growth of wild-type cells significantly, was added (Figure S6).

### 4.8 Fluorescence characterization for *pcnB* mutants

P1, P2 and wild-type strain harboring the pRFP were cultured in M9 medium. Fluorescence was measured (excitation, 587□nm; emission, 617□nm) in exponential growth phase and normalized to the culture OD_600_ value. Fluorescence intensity was calculated by comparing the relative fluorescence with respect to the wild-type strain harboring pRFP.

### 4.9 Statistical analysis

Error bars indicate SDs from 3 parallel experiments. Plots were generated in GraphPad Prism 8. Genome plots were generated in R using the OmicCircos package. All statistical analyses were performed using edgeR or Microsoft Excel 2016 (Microsoft Corporation).

## Abbreviations

DSB: double-strand break
SNV: single nucleotide variant
CBE: cytosine base editor
UGI: uracil DNA glycosylase inhibitor
ORI: origin of replication
ABE: adenine base editor
FACS: fluorescence-activated cell sorting
FADS: fluorescence-activated droplet sorting

## Authorship contribution statement

C. Z., X. L. and Y. Y. conceived and designed this project. Y. Y., X. L. and S. L. performed the experiments. Y. Y. analyzed the data and wrote the initial manuscript draft. X. L., X. X. and C. Z. supervised the research and gave advice on the experiments and manuscript. All authors revised and approved the manuscript.

## Acknowledgments

The authors would like to thank associate Prof. Qiang Li from the Department of Chemical Engineering, Tsinghua University, for providing support for Bioscreen C MBR.

