## Supplemental Figures, Tables and Data for "Base Editor-Mediated Large-Scale Screening of Functional Mutations in Bacteria for Industrial Phenotypes": supplementary information_20230531.docx

**This PDF file includes:**

Figs. S1 to S6

Tables S1 to S4

References

**Other Supplementary Materials for this manuscript include the following:**

Data S1 to S4

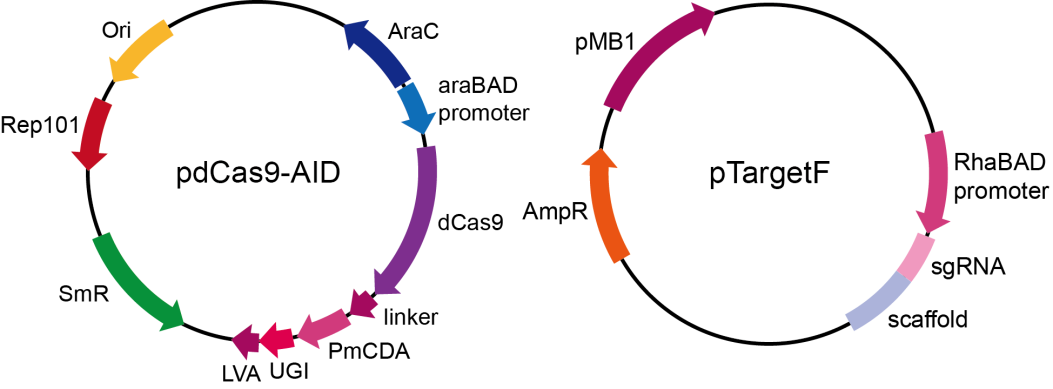

**Fig. S1.**

**The Target-AID plasmids for *E. coli*.** The plasmid pdCas9-AID contains the streptomycin resistant (SmR) gene and an arabinose inducible dCas9-PmCDA119-UGI-LVA fusion protein expression cassette.

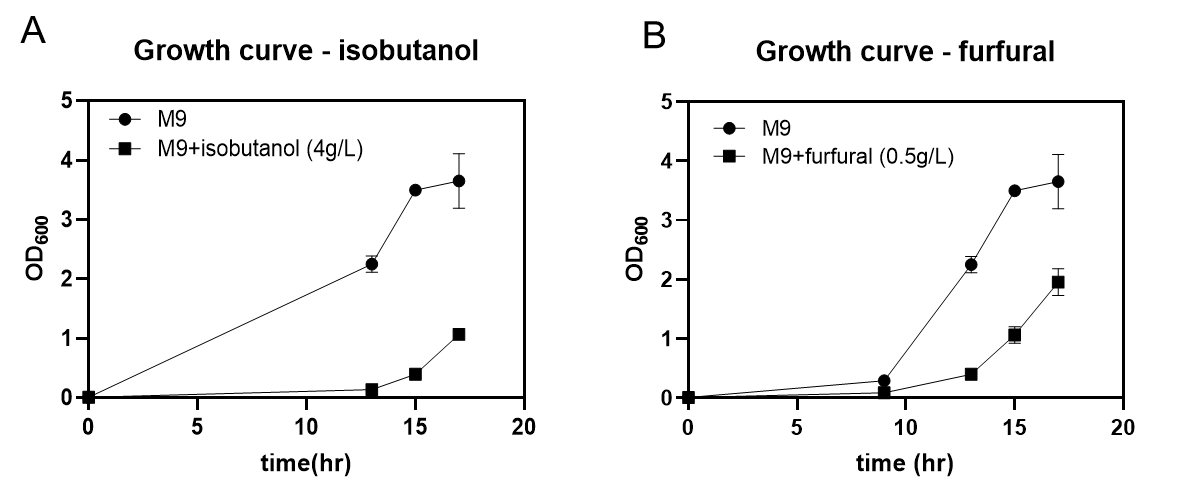

**Fig. S2.**

**Growth curve of *E. coli* BW25113 harboring non-targeting sgRNA in M9 medium containing** (A) 4 g/L isobutanol and (B) 0.5 g/L furfural. Strains were cultured in flasks. Results represent mean values and standard deviation of biological triplicates (N=3).

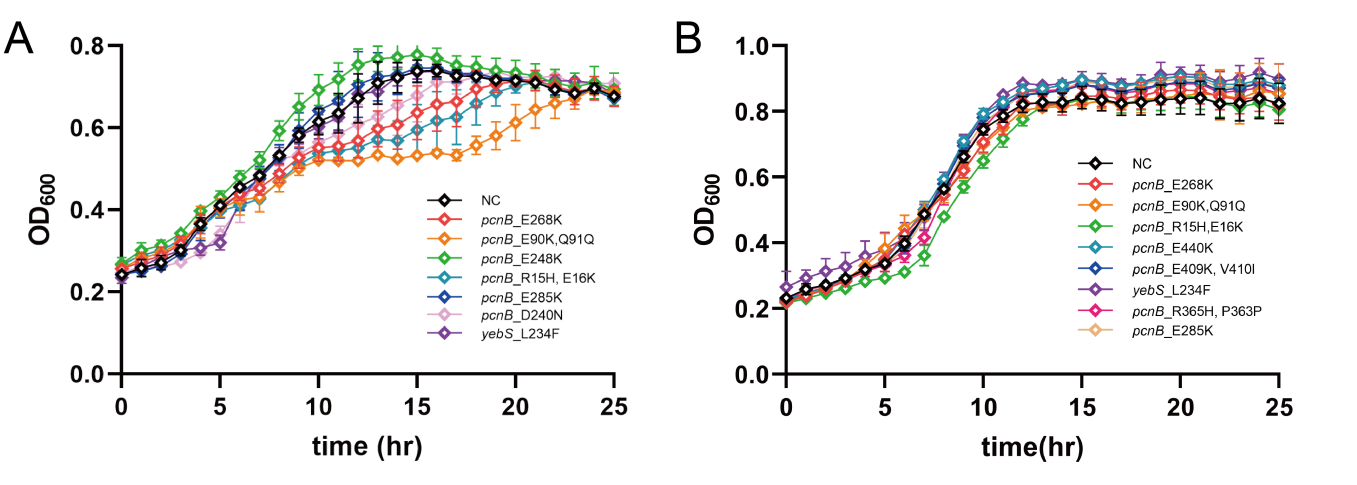

**Fig. S3.**

**Growth curves of the top 10 enriched variants in** (A) 4 g/L isobutanol and (B) 0.5 g/L furfural selections. The growth curves of the best mutants are shown in Fig. 3 in the main text. Results represent mean values and standard deviation of biological triplicates (N=3). NC indicates the strain harboring non-targeting sgRNA.

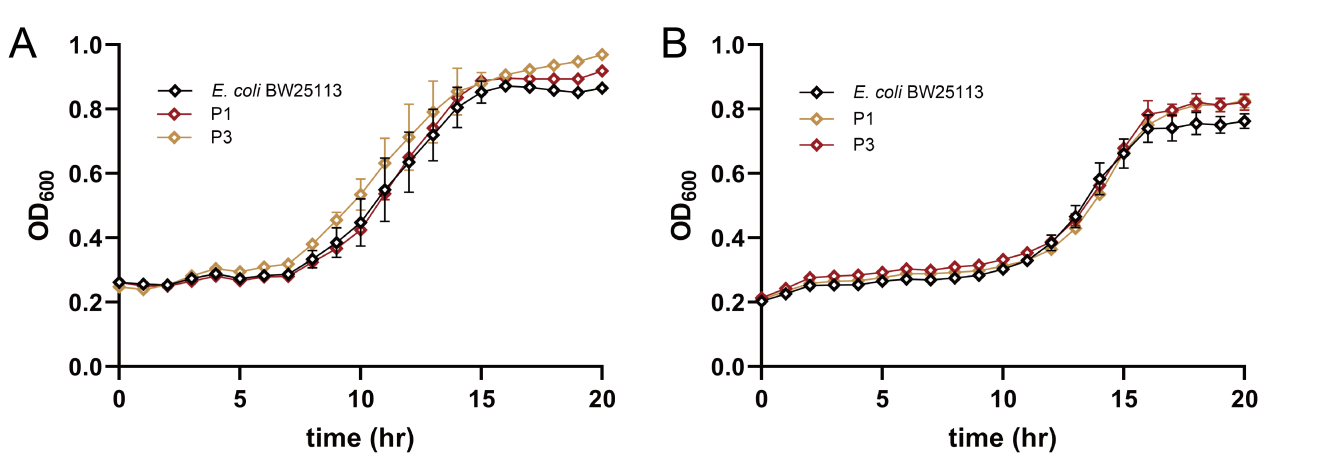

**Fig. S4.**

**Growth curves of P1, P3 and *E. coli* BW25113 in M9 medium containing (A) 5 g/L isobutanol and (B) 1 g/L furfural**. Results represent mean values and standard deviation of biological triplicates (N=3).

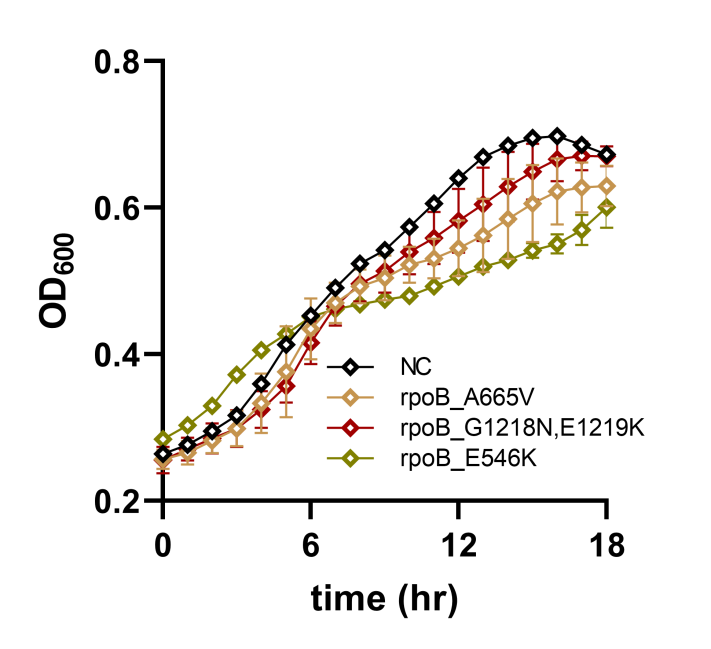

**Fig. S5.**

**Growth curves of** **rpoB_A665V,** **rpoB_G1218N,E1219K and rpoB_E546K in M9 medium containing 4 g/L isobutanol**. Results represent mean values and standard deviation of biological triplicates (N=3). NC indicates the strain harboring non-targeting sgRNA.

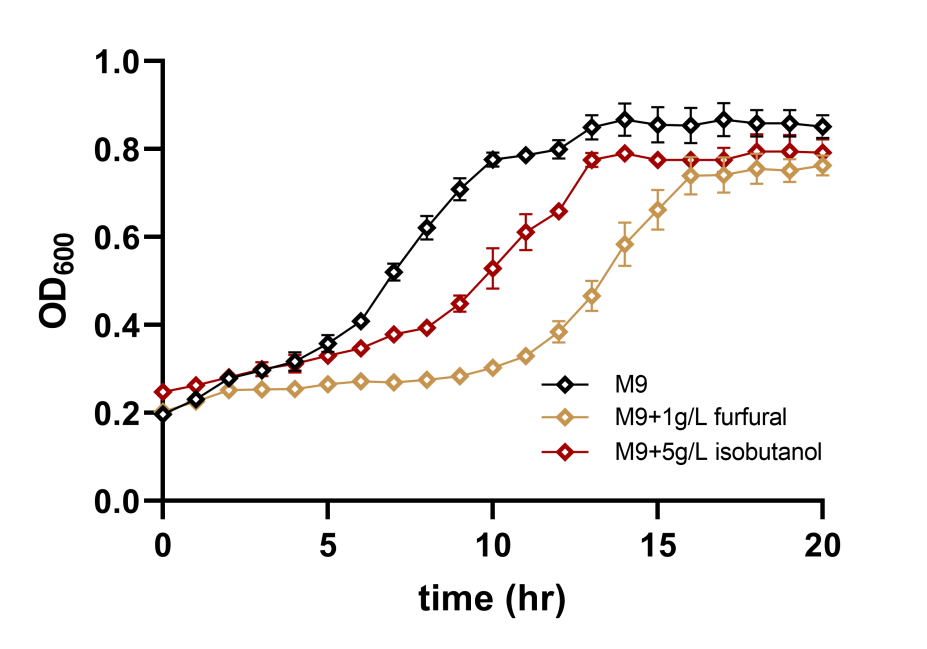

**Fig. S6.**

**Growth curve of** ***E. coli* BW25113 in M9 medium containing 5 g/L isobutanol or 1 g/L furfural**. Results represent mean values and standard deviation of biological triplicates (N=3). The strain was cultured in Bioscreen C MBR.

**Table S1. Editing efficiency of the sgRNA library.**

| Sample | Replicate1 | Replicate2 |
| --- | --- | --- |
| Editing efficiency | 83% | 82% |

**Table S2. Pearson coefficients for the screenings.**

| screening | Isobutanol | Furfural |
| --- | --- | --- |
| Pearson coefficient | 0.93 | 0.95 |

**Table S3. Strains and plasmids used in this work**

| Strains/plasmids | characteristics | sources |
| --- | --- | --- |
| Strains |  |  |
| *E. coli* DH5α |  | Biomed |
| *E. coli* BW25113 |  | Stored in our lab |
| *E. coli* s17-1 | Expresses sfGFP | ^1^ |
| Plasmids |  |  |
| pScI-dCas9-CDA-UL | dCas9-CDA-UGI-LVA expression cassette, CmR | Addgene #108551 |
| pSI-Target-AID-NG | Plasmid with nspCas9-NG sequence | Addgene #119861 |
| pdCas9-AID | dCas9-CDA-UGI-LVA with pBAD43, SmR | This study |
| pdCas9-AID-NG | pdCas9-AID with dead spCas9-NG, SmR | This study |
| pCas | Cas9 expression cassatte used in CRISPR/Cas9-based recombination, KmR |  |
| pTargetF | sgRNA expression plasmid used in CRISPR/Cas9-based recombination, SmR |  |
| pTarget | sgRNA expression plasmid with prhaBAD, pBR322, AmpR | This study |
| pTarget_rhaBAD_RFP | Vector for pTarget construction, carrying mCherry expression cassette, pBR322, AmpR | This study |
| pTarget-sfGFP36 | Expresses sgRNA targeting sfGFP, AmpR | This study |
| pTarget-sfGFP83 | Expresses sgRNA targeting sfGFP, AmpR | This study |
| pTarget-sfGFP166 | Expresses sgRNA targeting sfGFP, AmpR | This study |
| pTarget-sfGFP172 | Expresses sgRNA targeting sfGFP, AmpR | This study |
| pTarget-sfGFP408 | Expresses sgRNA targeting sfGFP, AmpR | This study |
| pTarget-NC | Expresses non-targeting sgRNA, AmpR | This study |
| pTarget-sgRNA8030 | Expresses sgRNA8030 targeting *yebS* in AID_library, used for enriched mutation reconstruction, AmpR | This study |
| pTarget-sgRNA20529 | Expresses sgRNA20529 targeting *pcnB* in AID_library, used for enriched mutation reconstruction, AmpR | This study |
| pTarget-sgRNA20521 | Expresses sgRNA20521 targeting *pcnB* in AID_library, used for enriched mutation reconstruction, AmpR | This study |
| pTarget-sgRNA20539 | Expresses sgRNA20539 targeting *pcnB* in AID_library, used for enriched mutation reconstruction, AmpR | This study |
| pTarget-sgRNA20542 | Expresses sgRNA20542 targeting *pcnB* in AID_library, used for enriched mutation reconstruction, AmpR | This study |
| pTarget-sgRNA20523 | Expresses sgRNA20523 targeting *pcnB* in AID_library, used for enriched mutation reconstruction, AmpR | This study |
| pTarget-sgRNA20548 | Expresses sgRNA20548 targeting *pcnB* in AID_library, used for enriched mutation reconstruction, AmpR | This study |
| pTarget-sgRNA20531 | Expresses sgRNA20531 targeting *pcnB* in AID_library, used for enriched mutation reconstruction, AmpR | This study |
| pTarget-sgRNA2520 | Expresses sgRNA20520 targeting *pcnB* in AID_library, used for enriched mutation reconstruction, AmpR | This study |
| pTarget-sgRNA20524 | Expresses sgRNA20524 targeting *pcnB* in AID_library, used for enriched mutation reconstruction, AmpR | This study |
| pTarget-sgRNA20510 | Expresses sgRNA20510 targeting *pcnB* in AID_library, used for enriched mutation reconstruction, AmpR | This study |
| pTarget-sgRNA20512 | Expresses sgRNA20512 targeting *pcnB* in AID_library, used for enriched mutation reconstruction, AmpR | This study |
| pTarget-sgRNA20516 | Expresses sgRNA20516 targeting *pcnB* in AID_library, used for enriched mutation reconstruction, AmpR | This study |
| pTarget-sgRNA26664 | Expresses sgRNA26664 targeting *gcvA* in AID_library, used for enriched mutation reconstruction, AmpR | This study |
| pTarget-sgRNA12223 | Expresses sgRNA12223 targeting *gcvA* in AID_library, used for enriched mutation reconstruction, AmpR | This study |
| pTarget-sgRNA12235 | Expresses sgRNA12235 targeting *gcvA* in AID_library, used for enriched mutation reconstruction, AmpR | This study |
| pTarget-sgRNA18177 | Expresses sgRNA18177 targeting *rpoB* in AID_library, used for enriched mutation reconstruction, AmpR | This study |
| pTarget-sgRNA18183 | Expresses sgRNA18183 targeting *rpoB* in AID_library, used for enriched mutation reconstruction, AmpR | This study |
| pTarget-sgRNA18195 | Expresses sgRNA18195 targeting *rpoB* in AID_library, used for enriched mutation reconstruction, AmpR | This study |
| pTarget-sgRNA30065 | Expresses sgRNA30065 targeting *rpoB* in AID_library, used for enriched mutation reconstruction, AmpR | This study |
| pTarget-sgRNA30122 | Expresses sgRNA30122 targeting *rpoB* in AID_library, used for enriched mutation reconstruction, AmpR | This study |
| pTargetF-P1 | sgRNA expression plasmid for creating mutation *pcnB*_R203H_E204K_D205N via CRISPR/Cas9-based recombination, SmR | This study |
| pTargetF-P3 | sgRNA expression plasmid for creating mutation *pcnB*_G183N via CRISPR/Cas9-based recombination, SmR | This study |
| pTargetF-gcvA-del | sgRNA expression plasmid for gcvA deletion via CRISPR/Cas9-based recombination, SmR | This study |
| pRFP | Expresses mCherry using Anderson promoter J23100 and RBS pFAB867, pBR322, AmpR | This study |

**Table S4. Primers and other oligonucleotides used in this work.**

| Primers/  oligonucleotides | Sequences (5’ to 3’) | Description |
| --- | --- | --- |
| Seq-F | GCGGCCTTTTTACGGTTC | Sequencing primer to confirm sgRNA sequences |
| pBAD-F | ttggctgttttggcggatga | Amplify pBAD43 vector |
| pBAD-R | gaattcctcctgctagcccaaaaa | Amplify pBAD43 vector |
| pdCas9-AID-F | tgggctagcaggaggaattcatggataagaaatactcaataggcttagCTATCG | Amplify dCas9-CDA-UGI-LVA expression cassatte |
| pdCas9-AID-R | tcatccgccaaaacagccaattatgcaaccagtccTAGCATCTTG | Amplify dCas9-CDA-UGI-LVA expression cassatte |
| dspCas9-NG-F1 | tgggctagcaggaggaattcatggacaagaagtactccattgggc | Amplify N-terminal of dspCas9-NG |
| dspCas9-NG-R1 | atccacgtcgtagtcggag | Amplify N-terminal of dspCas9-NG |
| dspCas9-NG-F2 | tctccgactacgacgtggatGCCatcgtgccccagtcttttct | Amplify C-terminal of dspCas9-NG |
| dspCas9-NG-R2 | CCTCCAGAACCTCCTCCACCgtcagccctgctgtctccac | Amplify C-terminal of dspCas9-NG |
| Target-AID-NG-backF | GGTGGAGGAGGTTCTGGAGG | Amplify plasmid backbone together with pBAD-R to construct pdCas9-AID-NG |
| NGS-seq-F | AGTAACGAGAAGGTCGCGTATTCA | NGS sequencing primer |
| NGS-seq-R | CTCGGTGCCACTTTTTCAAGTTGATAA | NGS sequencing primer |
| sgRNA-NC-F | tagtCTTCACTGTTCACACAATAG | Annealing to construct pTarget-NC by GoldenGate assembly |
| sgRNA-NC-R | aaacCTATTGTGTGAACAGTGAAG | Annealing to construct pTarget-NC by GoldenGate assembly |
| sgRNA-166-F | tagtGTCGTTACCAGAGTCGGCCA | Annealing to construct pTarget-sfGFP166 by GoldenGate assembly |
| sgRNA-166-R | aaacTGGCCGACTCTGGTAACGAC | Annealing to construct pTarget-sfGFP166 by GoldenGate assembly |
| sgRNA-172-F | tagtGTCAGCGTCGTTACCAGAGT | Annealing to construct pTarget-sfGFP172 by GoldenGate assembly |
| sgRNA-172-R | aaacACTCTGGTAACGACGCTGAC | Annealing to construct pTarget-sfGFP172 by GoldenGate assembly |
| sgRNA-408-F | tagtTGTATTCCAGCTTATGGCCC | Annealing to construct pTarget-sfGFP408 by GoldenGate assembly |
| sgRNA-408-R | aaacGGGCCATAAGCTGGAATACA | Annealing to construct pTarget-sfGFP408 by GoldenGate assembly |
| sgRNA8030-F | tagtTGTCTTGGGTGCCATTTTAC | Annealing to construct pTarget-sgRNA8030 by GoldenGate assembly |
| sgRNA8030-R | aaacGTAAAATGGCACCCAAGACA | Annealing to construct pTarget-sgRNA8030 by GoldenGate assembly |
| sgRNA20529-F | tagtCTTCACGGTAGCGCGTTTCC | Annealing to construct pTarget-sgRNA20529 by GoldenGate assembly |
| sgRNA20529-R | aaacGGAAACGCGCTACCGTGAAG | Annealing to construct pTarget-sgRNA20529 by GoldenGate assembly |
| sgRNA20521-F | tagtATATTCACACAACAGCTTAT | Annealing to construct pTarget-sgRNA20521 by GoldenGate assembly |
| sgRNA20521-R | aaacATAAGCTGTTGTGTGAATAT | Annealing to construct pTarget-sgRNA20521 by GoldenGate assembly |
| sgRNA20539-F | tagtGACCCACCAGGCGGCAGTTA | Annealing to construct pTarget-sgRNA20539 by GoldenGate assembly |
| sgRNA20539-R | aaacTAACTGCCGCCTGGTGGGTC | Annealing to construct pTarget-sgRNA20539 by GoldenGate assembly |
| sgRNA20542-F | tagtCTGCTCAGGCGTGGCGTTAG | Annealing to construct pTarget-sgRNA20542 by GoldenGate assembly |
| sgRNA20542-R | aaacCTAACGCCACGCCTGAGCAG | Annealing to construct pTarget-sgRNA20542 by GoldenGate assembly |
| sgRNA20523-F | tagtCTTCAAACAGGCGTGCCGGT | Annealing to construct pTarget-sgRNA20523 by GoldenGate assembly |
| sgRNA20523-R | aaacACCGGCACGCCTGTTTGAAG | Annealing to construct pTarget-sgRNA20523 by GoldenGate assembly |
| sgRNA20548-F | tagtCCTCGCGGCTTAGCACCTTG | Annealing to construct pTarget-sgRNA20548 by GoldenGate assembly |
| sgRNA20548-R | aaacCAAGGTGCTAAGCCGCGAGG | Annealing to construct pTarget-sgRNA20548 by GoldenGate assembly |
| sgRNA20531-F | tagtGCCGCCAACGTAATCACGGA | Annealing to construct pTarget-sgRNA20531 by GoldenGate assembly |
| sgRNA20531-R | aaacTCCGTGATTACGTTGGCGGC | Annealing to construct pTarget-sgRNA20531 by GoldenGate assembly |
| sgRNA20520-F | tagtTTCCGTGAAGTAGCGGGTAA | Annealing to construct pTarget-sgRNA20520by GoldenGate assembly |
| sgRNA20520-R | aaacTTACCCGCTACTTCACGGAA | Annealing to construct pTarget-sgRNA20520 by GoldenGate assembly |
| sgRNA20524-F | tagtTATCGTTCAGCAGGGTAGCG | Annealing to construct pTarget-sgRNA20524 by GoldenGate assembly |
| sgRNA20524-R | aaacCGCTACCCTGCTGAACGATA | Annealing to construct pTarget-sgRNA20524 by GoldenGate assembly |
| sgRNA20510-F | tagtGCTCGTTGAGCATCCCTTTT | Annealing to construct pTarget-sgRNA20510 by GoldenGate assembly |
| sgRNA20510-R | aaacAAAAGGGATGCTCAACGAGC | Annealing to construct pTarget-sgRNA20510 by GoldenGate assembly |
| sgRNA20512-F | tagtCTTCAGCTCGCAAGGCCAAC | Annealing to construct pTarget-sgRNA20512 by GoldenGate assembly |
| sgRNA20512-R | aaacGTTGGCCTTGCGAGCTGAAG | Annealing to construct pTarget-sgRNA20512 by GoldenGate assembly |
| sgRNA20516-F | tagtGTTTCGGGATTGCCAGTGAA | Annealing to construct pTarget-sgRNA20516 by GoldenGate assembly |
| sgRNA20516-R | aaacTTCACTGGCAATCCCGAAAC | Annealing to construct pTarget-sgRNA20516 by GoldenGate assembly |
| sgRNA26664-F | tagtAACCAATGAATGGCGAAACT | Annealing to construct pTarget-sgRNA26664 by GoldenGate assembly |
| sgRNA26664-R | aaacAGTTTCGCCATTCATTGGTT | Annealing to construct pTarget-sgRNA26664 by GoldenGate assembly |
| sgRNA12223-F | tagtGCAGACATATACCCGACAGT | Annealing to construct pTarget-sgRNA12223 by GoldenGate assembly |
| sgRNA12223-R | aaacACTGTCGGGTATATGTCTGC | Annealing to construct pTarget-sgRNA12223 by GoldenGate assembly |
| sgRNA12235-F | tagtGCGCAATCTGAAATCGAGGC | Annealing to construct pTarget-sgRNA12235 by GoldenGate assembly |
| sgRNA12235-R | aaacGCCTCGATTTCAGATTGCGC | Annealing to construct pTarget-sgRNA12235 by GoldenGate assembly |
| sgRNA18177-F | tagtAACTCCAACTTGGATGAAGA | Annealing to construct pTarget-sgRNA18177 by GoldenGate assembly |
| sgRNA18177-R | aaacTCTTCATCCAAGTTGGAGTT | Annealing to construct pTarget-sgRNA18177 by GoldenGate assembly |
| sgRNA18183-F | tagtTGCGTCCCTGATCCCGTTCC | Annealing to construct pTarget-sgRNA18183 by GoldenGate assembly |
| sgRNA18183-R | aaacGGAACGGGATCAGGGACGCA | Annealing to construct pTarget-sgRNA18183 by GoldenGate assembly |
| sgRNA18195-F | tagtGTCCACCGACCTCGGTGAAC | Annealing to construct pTarget-sgRNA18195 by GoldenGate assembly |
| sgRNA18195-R | aaacGTTCACCGAGGTCGGTGGAC | Annealing to construct pTarget-sgRNA18195 by GoldenGate assembly |
| sgRNA30065-F | tagtCACCAGTGCGACCATCGTAC | Annealing to construct pTarget-sgRNA30065 by GoldenGate assembly |
| sgRNA30065-R | aaacGTACGATGGTCGCACTGGTG | Annealing to construct pTarget-sgRNA30065 by GoldenGate assembly |
| sgRNA30122-F | tagtCTTCGAAGCCTGCACGTTCA | Annealing to construct pTarget-sgRNA30122 by GoldenGate assembly |
| sgRNA30122-R | aaacTGAACGTGCAGGCTTCGAAG | Annealing to construct pTarget-sgRNA30122 by GoldenGate assembly |
| P1-up-F | cgggacatacgcaactgca | Amplify the pcnB sequence to construct donor DNA for P1 |
| P1-up-R | ggaaacTcgTtaccAtAaaAatccggtacgtatgctgcg | Amplify the pcnB sequence to construct donor DNA for P1 |
| P1-down-F | cggatTttTaTggtaAcgAgtttccgggttaccaatcagacggataacg | Amplify the pcnB sequence to construct donor DNA for P1 |
| P1-down-R | aatttttgccgcaaggtgct | Amplify the pcnB sequence to construct donor DNA for P1 |
| P3-up-F | gtagcgggtaatggtcggga | Amplify the pcnB sequence to construct donor DNA for P3 |
| P3-up-R | CaactgTcgcctggtggAtcAccgtttccgtctggctcat | Amplify the pcnB sequence to construct donor DNA for P3 |
| P3-down-F | gaTccaccaggcgAcagttGcggaacagtttgcgcacc | Amplify the pcnB sequence to construct donor DNA for P3 |
| P3-down-R | gcccggtactaatcgcgg | Amplify the pcnB sequence to construct donor DNA for P3 |
| gcvA-up-F | gcagtagcggcgaacacac | Amplify the gcvA sequence to construct donor DNA for gcvA deletion |
| gcvA-up-R | ccgctaaatgccttacgagtttttgatgccgcagcacg | Amplify the gcvA sequence to construct donor DNA for gcvA deletion |
| gcvA-down-F | actcgtaaggcatttagcTgTtggtaatcgtttagacatggctattaaactttg | Amplify the gcvA sequence to construct donor DNA for gcvA deletion |
| gcvA-down-R | caccaatgacggacgtacca | Amplify the gcvA sequence to construct donor DNA for gcvA deletion |
| sgRNA-R | actagtattatacctaggactgagctagctgtcaag | Construction of pTargetF series plasmid |
| sgRNA-P1-F | TCCTAGGTATAATACTAGTCTTCACGGTAGCGCGTTTCCGTTTTAGAGCTAGAAATAGC | Construction of pTargetF-P1 plasmid together with sgRNA-R |
| sgRNA-P3-F | TCCTAGGTATAATACTAGTGACCCACCAGGCGGCAGTTAGTTTTAGAGCTAGAAATAGC | Construction of pTargetF-P3 plasmid together with sgRNA-R |
| sgRNA-gcvA-F | TCCTAGGTATAATACTAGTGACCCACCAGGCGGCAGTTAGTTTTAGAGCTAGAAATAGC | Construction of pTargetF-gcvA-del plasmid together with sgRNA-R |
| RFP-F | GTACAGTGCTAGCTTAATATCTTAATCTAGCTGGGGACTGTTTatggcttcctccgaagacg | Amplify RFP expression cassatte from pTarget_rhaBAD_RFP to Construct pRFP plasmid |
| RFP-R | actagtcGAGACGctctagtagaga | Amplify RFP expression cassatte from pTarget_rhaBAD_RFP to Construct pRFP plasmid |
| RFP-backbone-F | ctctctactagagCGTCTCgactagtGAATTCTCTAGAGTCGACCTGCAG | Amplify vector from pTarget_rhaBAD_RFP to Construct pRFP plasmid |
| RFP-backbone-R | GATATTAAGCTAGCACTGTACCTAGGACTGAGCTAGCCGTCAAGTAAATCCGTTCGTCTGCAATTATGG | Amplify vector from pTarget_rhaBAD_RFP to Construct pRFP plasmid |

**Data S1. (Separate file)**

Data S1 sgRNA_Library.xlsx

**Data S2. (Separate file)**

Data S2_mutation prediction in AID library.xlsx

**Data S3. (Separate file)**

Data S3_read counts in screening.xlsx

**Data S4. (Separate file)**

Data S4_fitness in screening.xlsx
